# Disorder Atlas: web-based software for the proteome-based interpretation of intrinsic disorder predictions

**DOI:** 10.1101/060699

**Authors:** Michael Vincent, Santiago Schnell

## Abstract

Intrinsically disordered proteins lack a stable three-dimensional structure under physiological conditions. While this property has gained considerable interest within the past two decades, disorder poses substantial challenges to experimental characterization efforts. In effect, numerous computational tools have been developed to predict disorder from primary sequences, however, interpreting the output of these algorithms remains a challenge. To begin to bridge this gap, we present Disorder Atlas, web-based software that facilitates the interpretation of intrinsic disorder predictions using proteome-based descriptive statistics. This service is also equipped to facilitate large-scale systematic exploratory searches for proteins encompassing disorder features of interest, and further allows users to browse the prevalence of multiple disorder features at the proteome level. As a result, Disorder Atlas provides a user-friendly tool that places algorithm-generated disorder predictions in the context of the proteome, thereby providing an instrument to compare the results of a query protein against predictions made for an entire population. Disorder Atlas currently supports ten eukaryotic proteomes and is freely available for non-commercial users at http://www.disorderatlas.org.

## Background

Intrinsically disordered proteins and protein regions do not form persistent secondary and tertiary structure under physiological conditions. A number of algorithms capable of predicting disorder in protein sequences play a critical role in disorder characterization efforts (Atkins et al., 2015; Monastyrskyy et al., 2014). A detailed treatment of the different disorder prediction methods is provided by Meng et al. (Meng et al., 2017). In this process, a user typically takes the sequence of a protein of interest and passes it as input into a selected disorder prediction algorithm, and residue-by-residue disorder predictions are commonly returned. These disorder scores are compared to a threshold value of disorder propensity to classify residues as being either disordered or ordered. Binary classifications are then used to compute the percent disorder, and length and location of continuous stretches of disordered residues. But what do these predictions mean? What is a significant disorder feature? And how can they be interpreted objectively? We put forth that standards for comparison are required to objectively interpret disorder predictions generated for a query protein. Disorder Atlas is a tool that enables users to utilize disorder predictions for full proteomes as standards for comparison. The purpose of the present article is to formally introduce this tool.

Disorder Atlas is web-based software that facilitates the interpretation of disorder predictions from amino acid sequence by comparing them with descriptive statistics, specific to both proteomes and disorder prediction tools, for identifying anomalous disorder features with respect to whole proteomic populations. We adopt the noun “atlas” under Gerardus Mercator’s original neologism of a descriptive reference rather than an extensive or complete collection. In Disorder Atlas, the reference is a set of proteome-based guidelines for standardizing intrinsic disorder predictions (Vincent et al., 2016), which are analogous to clinical guidelines used to evaluate whether an individual is overweight based on the body mass index distribution in the population. Although these guidelines do not provide a functional role of the disorder predictions, they provide a means of standardizing the prevalence of disorder in a query protein with respect to its proteome.

## Implementation

Disorder Atlas is primarily written in Python 2.7.12 and utilizes the Django model-view-controller framework. Disorder Atlas also interfaces with algorithms written in C and graphics modules written in JavaScript. All database operations are carried out using the Structured Query Language (SQL).

Pre-calculated proteome statistics are pulled from a PostgreSQL database, which have been computed from ten representative eukaryotic protein populations with minimal sequence redundancy and uncertainty (Vincent and Schnell, 2016; Vincent et al., 2016). Specifically, Disorder Atlas includes the *Saccharomyces cerevisiae, Dictyostelium discoideum, Chlamydonmonas reinhardtii, Drosophila melanogaster*, *Caenorhabditis elegans, Arabidopsis thaliana, Danio rerio, Mus musculus, Homo sapiens*, and *Zea mays* proteomes. However, Disorder Atlas accepts input sequences from any organism.

Reference proteome files obtained from UniProt served as the source of all protein sequence information and were subjected to multiple levels of filtering prior to disorder analysis. Disorder Atlas defines a sequence to be eligible for analysis if it does not include ambiguous and uncertain amino acid residues (such as B, J, O, U, X, and Z), and excludes sequences containing these features due to the variability observed in disorder prediction algorithms in processing input sequences containing these residue types (Vincent and Schnell, 2016; Vincent et al., 2016). The rationale behind this exclusion is that algorithmic variability for ambiguous and uncertain amino acids introduces error jeopardizing the accuracy of the analysis and standardization. Furthermore, redundancy was minimized by using UniRef100 reference cluster information obtained from the UniProt identification mapping service (Suzek et al., 2007; UniProt, 2015). The redundancy minimization procedure has been described in detail elsewhere (Vincent et al., 2016).

Currently, Disorder Atlas supports IUPred (Dosztanyi et al., 2005a; Dosztanyi et al., 2005b) and DisEMBL (Linding et al., 2003) disorder predictions (as well as consensus agreement between the two). Briefly, IUPred predicts disorder by assessing pairwise interresidue interaction energy and identifies disordered residues as those that are incapable of forming stabilizing interresidue interactions (Dosztanyi et al., 2005a; Dosztanyi et al., 2005b). DisEMBL is a suite of three neural networks algorithms that predict the presence of disorder: COILS, HOTLOOPS, and REM465, referred to by Disorder Atlas as DisEMBL-C, DisEMBL-H, and DisEMBL-R, respectively. Each describes disorder as a two-state model, classifying residues as either disordered or ordered (Linding et al., 2003). DisEMBL-C utilizes secondary structure prediction to assign disorder/order classifications, and classifies residues as disordered if they are present in loops (Linding et al., 2003). Residues are predicted to belong to loops if they do not belong to either alpha-helices, 310-helices, or beta-strands (assumes all residues belonging to loops are disordered) (Linding et al., 2003). Building on the reasoning that not all loops are disordered but all disordered residues are found within loops, DisEMBL-C classifications represent an overestimate of disorder. For improved disorder classifications, Linding et al. (Linding et al., 2003), implemented DisEMBL-H, which classifies residues (contained within loops) as disordered only if its alpha-carbon has a high temperature factor (B factor). Lastly, the DisEMBL-R neural network has been trained using non-assigned electron densities from X-ray crystallography (XRC) data contained within the Protein Data Bank, and assumes residues with missing XRC coordinates (as defined by REMARK 465 entries in PDB files) are disordered (Linding et al., 2003). The threshold values suggested by the algorithm developers are used to make algorithm-specific residue-by-residue disorder classifications. The consensus predictions are merely computed from a binary order/disorder residue-by-residue classification that is defined by a combination of prediction algorithms. This is a simple, transparent consensus prediction. If a specified combination of prediction algorithms agree that a residue is disordered, then that residue is classified as disordered.

Disorder Atlas also provides information about protein hydropathy, charge distribution and other disorder-relevant parameters as predicted by CIDER (Holehouse et al., 2017). In addition to its single protein assessment features, Disorder Atlas also provides tools for browsing disorder at the proteome-level and for conducting an exploratory search for proteins with disorder features of interest. These tools are described in detail in the next section.

## Results and Discussion

The Disorder Atlas web-based interface provides access to three tools for interpreting disorder predictions: (1) a proteome-level disorder browser, (2) an individual protein analysis tool, and (3) a proteome exploratory search tool. Each tool is based on descriptive population statistics, and presents the standing of disorder features in relation to a proteomic population based on quantitative guidelines for standardizing disorder predictions (Vincent et al., 2016). Disorder Atlas utilizes two physicochemical-based disorder prediction algorithms, IUPred-L and DisEMBL. Consensus disorder predictions are also presented, which provide more conservative disorder annotations.

### The Proteome Browser

This tool provides the distribution of three disorder statistics at the proteome level: (1) the disorder content, (2) the longest continuous disorder region (CD_L_), and (3) the longest CD_L_ percentage of length (LCPL). The disorder content is simply the percentage of disordered residues contained within a protein sequence. The CDL is the longest continuously disordered region in a protein, defined using the theoretical minimum of two consecutive disordered residues (while a CD segment of two amino acids may be structurally unimportant, it includes all possible predicted CD segments and avoids using a subjective minimum length that could potentially exclude valid short CD regions). The LCPL defines the percentage of the total protein length accounted for by the CDL and is useful for identifying a statistically relevant long CD segment in proteins having a primary sequence length exceeding previously reported protein length thresholds (Vincent et al., 2016). For each of these disorder statistics, users can access the proteome browser to visualize statistical distributions, percentiles, and expected values.

### Individual Protein Analysis Tool

For single protein analyses, users can provide either the (1) UniProt accession number, or (2) FASTA sequence, and the name of the proteome to which the sequence belongs. Following submission, the disorder propensity, as well as the standing of the disorder content, CD_L_, and LCPL with respect to the proteome, is presented (two sample result plots are displayed in **Figure 1A**). Additionally, Disorder Atlas also presents protein hydropathy and charge distribution and other disorder-relevant parameters generated by localCIDER version 0.1.7 (Holehouse et al., 2017). The generated histograms, boxplots, and disorder propensity charts can be downloaded as either a PNG or SVG file. All pages can be easily saved as a PDF file from any web browser print menu.

**Figure 1.**
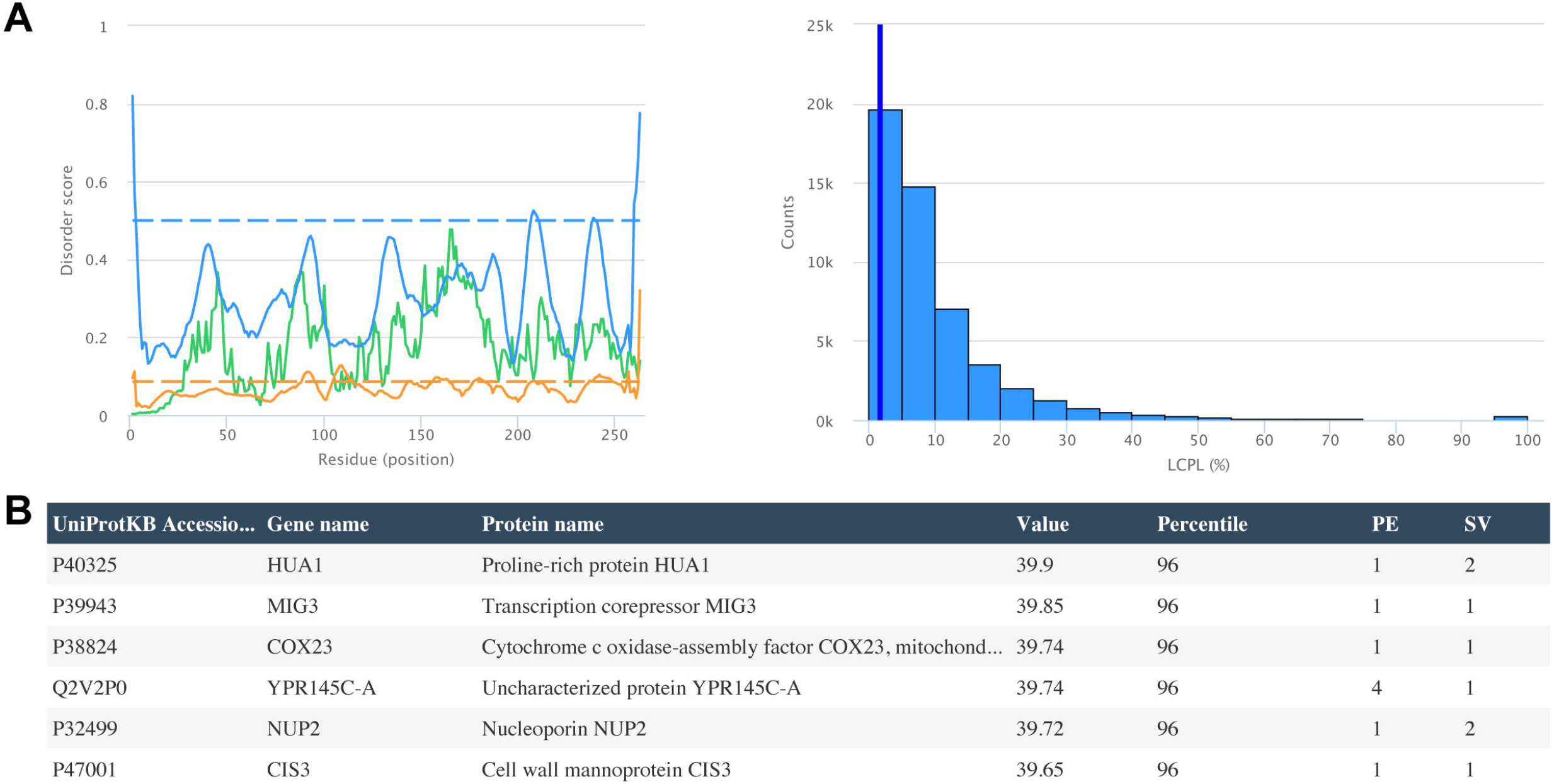
**(A)** Example output from the individual protein analysis tool. The disorder propensity predicted by IUPred-L (green), DisEMBL-H (orange), and DisEMBL-R (blue) for Chymotrypsinogen B (P17538) is shown on the left, whereas its LCPL predicted by DisEMBL-R (blue vertical line) is shown together with the *Homo sapiens* DisEMBL-R LCPL distribution on the right. **(B)** Example output from the exploratory search tool. A search was conducted to find *Saccharomyces cerevisiae* proteins with a DisEMBL-R-predicted disorder percentage of less than 40%. A truncated table displaying the six most disordered proteins meeting the search criteria is shown.

### Proteome Exploratory Search Tool

Disorder Atlas can provide an exploratory search for proteins with a disorder feature of interest. To conduct this search, users specify the proteome to be searched, the disorder metric and prediction method of interest, and whether they would like to conduct a value-based or percentile-based search. For example, a user could search for all *Saccharomyces cerevisiae* proteins with a DisEMBL-R-predicted percent disorder of less than 40% (a truncated result table for this search is displayed in **Figure 1B**), or they could look for all *Homo sapiens* proteins with a CD_L_ above the 75^th^ percentile (not shown). After submitting a search query Disorder Atlas returns a list of proteins meeting the specified criteria together with their associated statistical values. The search results can be exported in a variety of file formats, including CSV, JSON, PDF, SQL, TXT, and XML.

### Improvements over existing protein disorder resources

A handful of databases exist that contain disorder annotations, including D2P2 (Oates et al., 2013), DisProt (Piovesan et al., 2017), and MobiDB (Di Domenico et al., 2012; Potenza et al., 2015). These resources provide users with valuable information concerning protein regions predicted to be disordered, and provides experimental support of disorder predictions when available. Importantly, the number of protein sequences having experimental information relevant to protein disorder represent only a small fraction of the total number of known sequences. As such, researchers are often confronted with the problem of interpreting computational disorder predictions without this information.

Intrinsic disorder is a prevalent feature of many proteomes (Vincent and Schnell, 2016; Vincent et al., 2016), and such, the mere prediction of disorder (to any degree) is insignificant. In cases where experimental information regarding the disorder content for a protein of interest is unavailable, software is needed to enable a standardized interpretation of these predictions. For example, if a given protein is found to contain 20 amino acid long disordered region, is this significant at the proteome-level? Or if 30% of the residues in a protein are predicted to be disordered, what should be made of this prediction from the proteomic vantage point? The aforementioned databases cannot answer these questions when experimental information is unavailable. Disorder Atlas addresses a major need that is unmet by any existing resources containing disorder annotations: it provides researchers with a tool for standardizing disorder predictions with respect to whole proteomes. More formally, Disorder Atlas provides standards for comparison that can be used to compare the disorder content of a query protein against predictions made for the rest of the proteome of origin. This software provides a simple, accessible solution to the non-trivial problem of interpreting the meaning of results generated by disorder prediction algorithms.

## Conclusions

Numerous intrinsic disorder prediction algorithms exist and as our understanding of disorder expands, more algorithms are developed that incorporate different definitions of disorder. While the number of tools that predict intrinsic disorder expands, the development of separate tools and guidelines needed to objectively interpret the meaning of these disorder predictions lags considerably behind. The absence of guidelines for standardizing the prevalence of disorder restricts the interpretation of the algorithm-generated predictions by scientists without a sophisticated understanding of structural biology and protein informatics. Disorder Atlas is web-based software that aims to bridge this gap by providing accessible and versatile tools for standardizing protein disorder predictions. The guidelines associated to Disorder Atlas provide a meaningful improvement in the methods of reporting protein disorder.

We plan to support additional proteomes within the upcoming year after implementing an automated disorder prediction and analysis pipeline. Following implementation of this system, users will have access to a multitude of prokaryotic and eukaryotic proteomes. We further envision that additional disorder prediction algorithms will be supported in the future as well.

## Declarations

### Availability of data and material

The protein sequences, their disorder predictions, hydropathy, charge distribution and other disorder-relevant parameters that support Disorder Atlas are available in Dryad with the identifier DOI 10.5061/dryad.sm107. (Vincent and Schnell, 2016)

Disorder Atlas is freely available for non-commercial users at http://www.disorderatlas.org. The computer code is published with a dual license scheme (GNU General Public License and a modified version of the Limited GNU Public License) to permit use and further development by non-profit and commercial entities.

## Acknowledgements

We thank Dr. Arash S. Soleimanpour (University of Michigan) for testing Disorder Atlas.

